# Chemotaxis model for human breast cancer cells based on signal-to-noise ratio

**DOI:** 10.1101/292300

**Authors:** S. Lim, H. Nam, J.S. Jeon

## Abstract

Chemotaxis, a biased migration of cells under a chemical gradient, plays a significant role in diverse biological phenomena including cancer metastasis. Stromal cells release signaling proteins to induce chemotaxis, which further causes organ-specific metastasis. Epidermal growth factor (EGF) is an example of the chemical attractants, and its gradient stimulates metastasis of breast cancer cells. Hence, the interactions between EGF and breast cancer cells have long been a subject of interest for oncologists and clinicians. However, most current approaches do not systematically separate the effects of gradient and absolute concentration of EGF on chemotaxis of breast cancer cells. In this work, we develop a theoretical model based on signal-to-noise ratio to represent stochastic properties and report our microfluidic experiments to verify the analytical predictions from the model. The results demonstrate that even under the same EGF concentration gradients, breast cancer cells can reveal distinct chemotaxis patterns at different absolute concentrations. Moreover, we found that addition of EGF receptor antibody can promote chemotaxis at a low EGF level. This apparently counterintuitive finding suggests that EGF receptor-targeted therapy may stimulate metastasis of breast cancer at a particular condition, which should be considered in anticancer drug design.

## INTRODUCTION

Cells have biased directionality in response to external chemical attractants (1). Known as chemotaxis, it is related to diverse physiological and pathological phenomena (2–5), such as wound healing (2) and cancer metastasis (5). In case of cancer metastasis, organ-specific stromal cells release signaling proteins that induce chemotaxis and attract cancer cells, causing organ-specific metastasis (5, 6). For example, epidermal growth factor (EGF) promotes invasion of breast cancer cells (6), and fractalkine stimulates metastasis of lung cancer (1). Notwithstanding the clinical significance, the detailed process of chemotactic response related to metastasis has yet to be clarified.

Previous studies on eukaryotic chemotaxis model provide a guide to understanding chemotaxis of cancer cells since cancer cells belong to a group of eukaryotic cells. Eukaryotic cells employ an asymmetrical cellular distribution of substrate-bound receptors to recognize a chemical gradient (7). The establishment of the asymmetric distribution involves random errors that originate from thermal fluctuations of local ligand concentrations (8). These errors exist in both binding reactions (8–11) and signal transduction reactions (12, 13). However, most of the previous studies considered the extraneous noise negligible (13–17). We speculated that the external noise from thermal fluctuations can be significant, especially when chemical concentrations are low (18), and adopted a signal-to-noise ratio (SNR) as a physical index to investigate the signal-noise relationship (8, 13–17, 19–21). While there have been efforts to understand chemotaxis using SNR, many of the previous studies investigated chemotaxis of an amoeba, *Dictyostelium discoideum* (13–15, 20, 21), as a simple eukaryotic model organism with directional movement. Since amoebae exhibit different characteristics from mammalian cells (1, 22), the transferability of the results to mammalian cells may be limited. Also, there have been efforts to model chemotaxis of leukocytes (8) and axons (16, 17) where SNR was used with Bayesian theory (16, 17), and a relation between SNR and the probability for chemotaxis was analyzed (8). However, to our knowledge, theoretical modeling with SNR approach in cancer cell chemotaxis has not yet been reported.

Experimentally, chemotaxis assays have been used to investigate chemotaxis of mammalian cells such as leukocytes (8–11, 23–25), endothelial cells (4), fibroblasts (26), and axons (16, 17, 27). The conventional platforms, including Zigmond chamber (23), Dunn chamber (24), and filter assay (28), have contributed to the development of chemotaxis models. However, they lack the ability to track cancer cell migration in a 3D environment with the precise maintenance of gradients and reconstruction of the microenvironment. Development of microfluidic system enables modeling of 3D tumor microenvironment with a linear gradient of ligand concentration and facilitation of real-time imaging (25, 29, 30). This setup is especially beneficial in investigating chemotaxis of tumor cells since the tumor microenvironment dramatically influences the cellular response to external stimuli (6) and the mass transfer of growth factors (31, 32). Some previous studies utilize the advantageous microfluidic platform for studying cancer cell chemotaxis (33–35), and we take a step further by combining SNR-based theoretical modeling with and microfluidic-based experimental analysis of cancer cell chemotaxis.

This study is primarily focused on chemotaxis of a human metastatic breast cancer cell, MDA-MB-231, under chemotactic EGF gradient. Using SNR, we have derived an analytical expression to depict cancer cell chemotaxis and validated the theoretical results by establishing a microfluidic system for tracking cancer cell chemotactic migration. This combined theoretical and experimental cancer cell chemotaxis model attempts to fill the gap of the previously unexplored area of SNR-based mammalian cell chemotaxis models. The results demonstrate that the absolute value of concentration governs chemotaxis under the same linear gradient and that the reduction in the number of EGF receptor (EGFR) under some conditions may promote chemotaxis at a low level of EGF.

## MATERIALS AND METHODS

### Cell culture

MDA-MB-231 cells (Korean Cell Line Bank) were cultured in Dulbecco’s modified Eagle medium (Lonza) supplemented with 10% fetal bovine serum (Gibco) and 1% antibiotic/antimycotic solution (Gibco), and kept in a CO_2_ incubator at 37°C and 5% CO_2_. Media were supplemented with human recombinant EGF (Sigma) at two different concentrations (150 ng/ml and 1 ng/ml) to induce linear EGF gradient for chemotaxis. EGFR antibody (Millipore) was added at two different concentrations (1 ng/ml and 10 μg/ml) to reduce the effective number of free receptors without genetic manipulation. The EGFR antibody was added to the medium channels before the experiment and incubated for 4 hours.

### Microfluidic system

A microfluidic device with two medium channels and one gel channel was fabricated with soft lithography using PDMS (Dow Corning). Plasma treatment bonded the PDMS layer to a slide glass, and the device was placed in a drying oven to recover hydrophobicity. Fabrication details were described previously (36). The basal thrombin solution, obtained by dissolving 20 μl thrombin (100 U/ml) in 500 μl cell culture medium, was used to suspend cancer cells at 150,000 cells/ml to generate cancer cell containing thrombin solution. Then, 10 μl fibrinogen solution (5 mg/ml) was mixed with 10 μl cancer cell containing thrombin solution, and the resulting fibrin gel is injected into the gel channel of the device. The devices were incubated in a humid chamber for 15 min in the incubator for gelation. Once hydrogel is formed, EGF free growth medium was introduced in two medium channels, and the devices were incubated for 1 day. When refreshing the media, EGF supplemented medium was introduced in one medium channel, and EGF free growth medium was in the other channel. In order to establish EGF gradient, a syringe pump was connected to the microfluidic system. The syringe pump extracted the medium in both medium channels through an outlet port at 100 μl/hr, and the solutions in the reservoirs replenished the chemicals continuously and maintained the chemical gradient across the device (Figs. 2*A* and S2).

**FIGURE 2.**
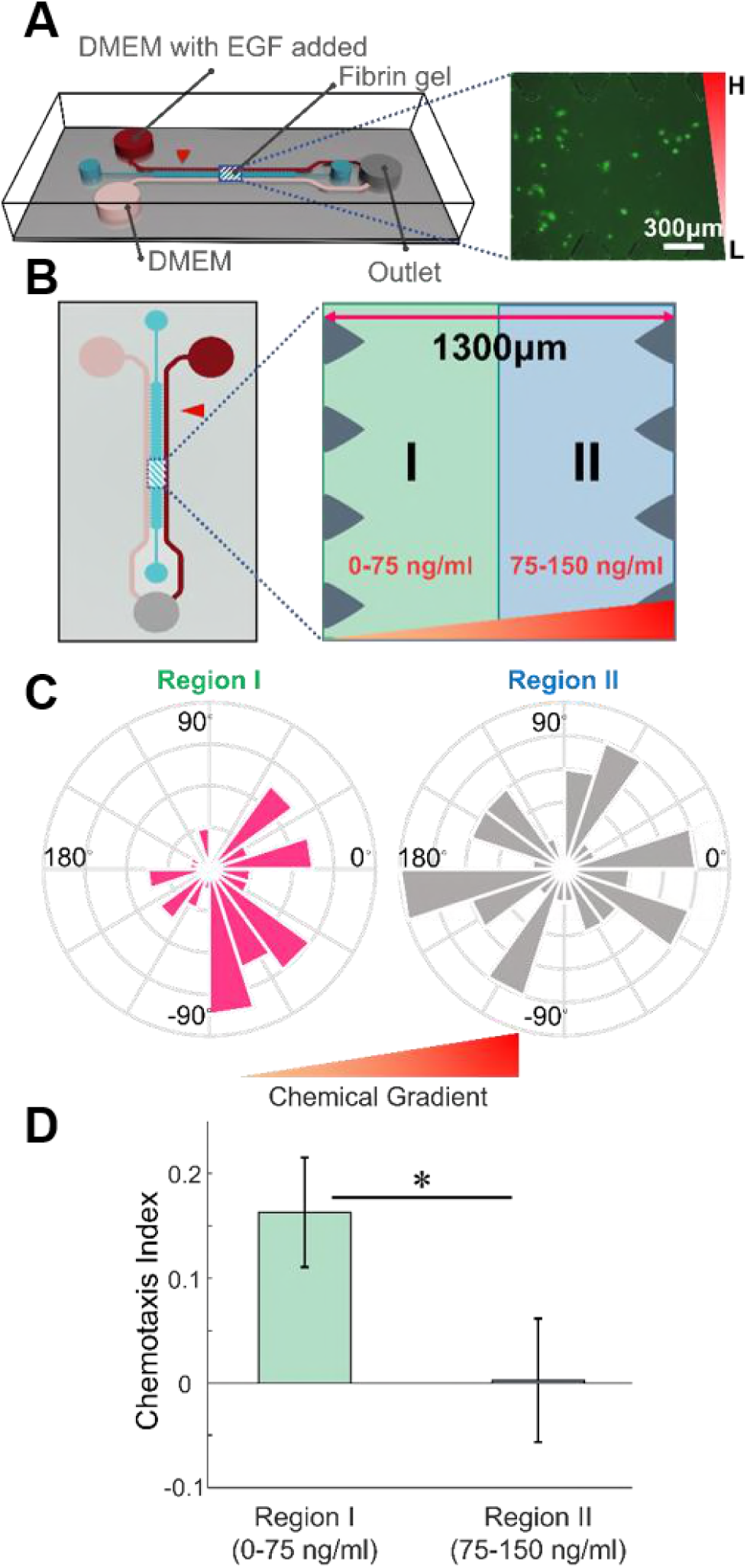
Chemotaxis of MDA-MB-231 in the microfluidic device. (A) A microfluidic device with center gel channel and two media channels allows formation and maintenance of an EGF chemical gradient in the center. Fluorescently-labeled breast cancer cells are placed in the center gel region and tracked for their migration behavior. The inverted right triangle represents the EGF gradient. (B) Top view of a microfluidic device and imaginary division of center gel channel for the cells according to their positions (region I in green, region II in blue) are shown. (C) Angle histograms and (D) CI values of the cells in the region I and region II show a more biased migration behavior for cancer cells in region I. Mean ± SEM (n = 50 for Region I and 51 for Region II). * p<0.05.

### Cell tracking and Statistical Analysis

Cancer cells were fluorescently stained with cell tracker at 5 μM for 45 min (Molecular Probes). Cells were photographed for 3 hours every 30 minutes using Observer Z1 (Zeiss) microscope, and images were preprocessed using AxioVision (Zeiss). The trajectory of cancer cells was tracked with the program of Piccinini et al. using MATLAB 9.2.0 (MathWorks) (37).

To quantify the extent of chemotaxis, we calculated chemotaxis index (CI), defined by a ratio of a net displacement toward chemical gradients to the total migration distance.

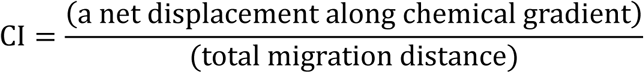

Also, to compare the magnitude of bias in cell orientations, from azimuth angles of cell orientations, we extracted a parameter κ by fitting a von Mises distribution to the angle data (38). The von Mises probability density function for the angle θ is a form of

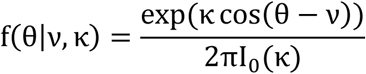

where I_0_(∙) is the modified Bessel function of order 0, ν is the mean of the distribution, and κ is an indicator of the data bias. The distribution of data is close to a uniform distribution as κ is close to zero.

Measurements are obtained from five to twelve independent devices for chemotaxis of linear gradient and chemotaxis with addition of EGFR antibody experiment, respectively. We performed the Kolmogorov–Smirnov test for normality, and did the two-tailed Wilcoxon rank sum test and independent t-test for group comparison. All statistical tests were executed with MATLAB 9.2.0.

### Simulation details

The chemotaxis simulation assumes a three-dimensional system of a cell using a sphere with a radius of 7.5 μm. The spherical cell consists of 720 segments divided by 20 latitudes and 36 longitudes, and each segment has the number of receptors proportional to the area of each segment. The total number of receptors across a cell is 800,000 molecules. Each cell has a direction vector, along which it moves by 0.4 μm at each simulation step (corresponding to 1 minute) for 180 steps (corresponding to 3 hours). The initial position of a cell is determined by uniform distribution in the region I or region II (Fig. 1*B*). The number of iterations for the chemotaxis simulation is 5,000 iterations in each region.

**FIGURE 1.**
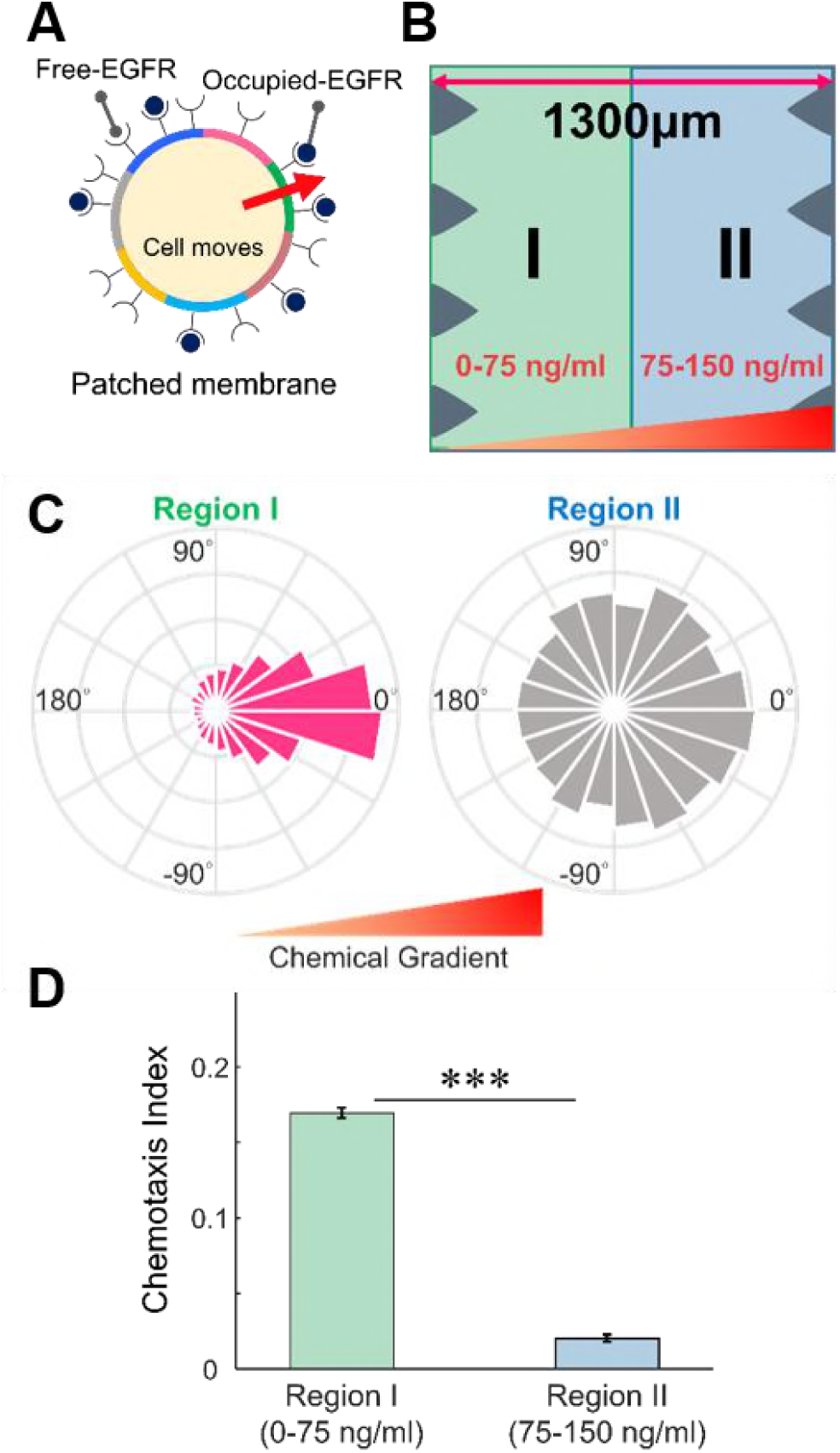
Chemotaxis simulations under equal chemical gradients and different concentrations. (A) Cancer cells are simulated to move towards the direction determined by the largest number of EGF bound receptors. (B) Cells are placed in 1300 μm wide channel with linear concentration gradient for the simulation. An imaginary border divides the channel according to positions of cells (region I in green, region II in blue). (C) Both the angle histograms and (D) chemotaxis index values of the cells placed in the region I and region II demonstrate that the tendency of chemotaxis decreases as the local EGF concentration increases while the EGF gradient remains the same. Mean ± SEM (n = 5000 for each region). *** p<0.001.

A direction vector is updated at the beginning of each step, based on the numbers of EGF-bound receptors of all segments. The number of EGF-bound receptors is randomly drawn from a normal distribution of mean and variance as derived in Eqs. (S11) and (S12) for each segment (Supporting Material). Since the addition of EGF to MDA-MB-231 forms an average of 24 filopodia (39), the cell selects 24 segments with the largest number of EGF-bound receptors. The direction vector is determined by the direction of a uniformly selected segment among the 24 segments.

## RESULTS

### Ligand concentration value governs chemotaxis under the same gradient

While chemotaxis occurs under a chemical gradient, the sole effect of local concentration on chemotaxis (separated from the gradient effect) is not always apparent. We simulated a chemotactic system (Fig. 1*A*) where cells freely move in a 1300 μm-wide channel under a linear concentration gradient of EGF (0 to 150 ng/ml). A cell determines its moving direction by calculating the distribution of EGF-bound receptors on its surface, and a local number of EGFs is allowed to fluctuate with a Gaussian noise (a detailed model description is given in the Supporting Material). To see the effect of initial positions (and consequent local EGF concentrations) of cancer cells, we virtually divided the space into two regions: region I with EGF concentration 0 to 75 ng/ml, and region II with 75 to 150 ng/ml (colored in green and blue, respectively, in Fig. 1*B*). While the local concentration of EGF varies in the two regions, the concentration gradient of the two regions remained the same. The simulated angle distribution of net cell migration implies that the cells at lower concentrations showed more biased movement than their counterparts at higher concentrations. (Fig. 1*C*). This result can be quantitatively verified by comparing the von Mises distribution parameter κ, which compares cellular orientation angles; the values of κ are 2.1882 and 0.3884 for region I and region II, respectively. We also investigated the chemotaxis index (CI) values, which represent a ratio of a net displacement toward chemical gradients to the total migration distance. As depicted in Fig. 1*C*, region I shows the CI of 0.166 ± 0.003, while CI for region II is 0.020 ± 0.001. Taken together, these results illustrate that chemotaxis of cells is less significant in a higher local EGF concentration, even under the same gradient, at least in the framework of the simulation.

### MDA-MB-231 exhibits distinct chemotaxis at low EGF concentration

In order to validate the simulation results, we conducted chemotaxis experiments in the microfluidic system (Fig. 2*A*). The concentration of EGF is 0 ng/ml at one media channel and 150 ng/ml at the other channel, and the concentrations are maintained by a syringe pump that generates a linear gradient of EGF across the device, which further allows us to calculate local EGF concentrations from horizontal positions. To see the effect of initial positions (and consequent local EGF concentrations) of cancer cells, we virtually divided the space into two regions as before; region I and II contained 50 cells, 51 cells, respectively. (Fig. 2*B*). The cellular positions had been recorded for 3 hours, and a chemotaxis pattern is similarly quantified by the azimuthal angle distributions of the cell migration directions and the CI values (Figs. 2*C* and 2*D*). As expected, the motion of cells in region II (higher local concentration) is closer to random motion whereas that in region I is more directed. (κ = 0.9013 for region I and 0.0443 for region II). We also observed the dramatic difference in CIs between two spatial groups of cells (0.163 ± 0.052 for region I and 0.002 ± 0.059 for region II). The two measures together indicate that local EGF concentration difference leads to a difference in chemotaxis behavior even under the same gradient, as predicted by the simulations. It is worth noting that the cell migration speed did not show a noticeable difference (0.405 ± 0.027 μm/min vs. 0.449 ± 0.026 μm/min, Fig. S3), which implies that the different chemical concentrations do not have a significant effect on the speed of cells, but only on directionality.

### SNR is a key parameter in the pattern of chemotaxis

Now the question of why the local concentration affects chemotaxis remains. In order to address the question, we developed a theoretical framework that incorporates concentration fluctuations in a biochemical reaction of receptors and ligands. The fluctuations can attenuate the signal generated from a biochemical process. We introduce a SNR-based chemotaxis model, where a cell decides its moving direction based on a conditional probability that it finds a correct direction given the concentration gradient between the two ends of the cell, with the additive white Gaussian noise model. As derived in Supporting Material, the conditional probability is given by

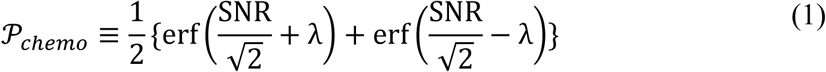

where erf(∙) is an error function, λ is a normalized threshold of decision making, and SNR is the signal-to-noise ratio, which is calculated as follows (see Supporting Material):

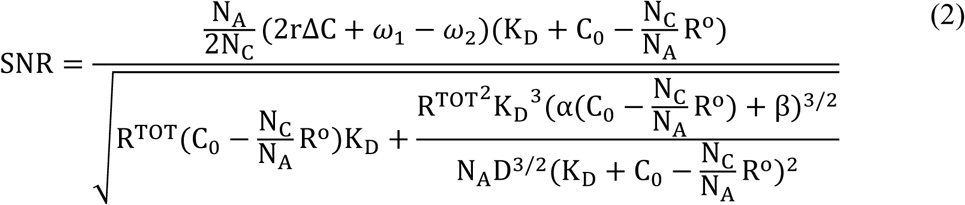

where ω_1_ and ω_2_ are respectively related to the numbers of bound receptors at posterior and anterior areas of a cell under the chemical gradient, according to the following expressions:

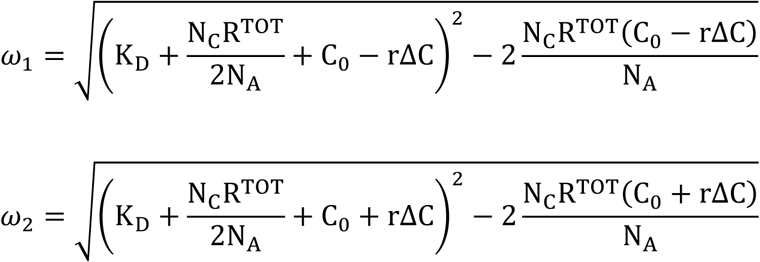

Here, C_0_ is the reference concentration of ligand, α is the binding rate, β is the dissociation rate, R^TOT^ is the total number of receptors, K^D^ is the dissociation constant, r is the half-length of a cell, ∆C is the gradient of ligand, N^A^ is Avogadro’s constant, N^C^ is the cell density, and D is the ligand diffusivity. We assume that the conditional probability serves a role of the chemotaxis propensity, and propose it to be an effective determinant of a chemotaxis pattern.

In previous works, chemotaxis has been modeled in terms of the difference in bound receptor numbers across a cell, which captures asymmetry of local concentrations around the given cell (33, 35). For a given number of bound receptors R°, the difference of R° at anterior and posterior areas of the cell, denoted as ∆R°, is given by the following formula (see Supporting Material for derivation):

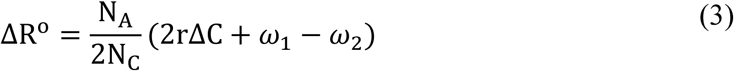

∆R° and 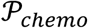 are both monotonically decreasing functions of local concentrations (Fig. 3, dark blue); this implies that while the concentration gradient is kept same, both degrees of chemotaxis decreases as the absolute concentration of the ligand increases, consistent with the simulation and experiment results reported in the previous sections. However, the two models predict different chemotaxis behaviors when EGFR concentrations are lowered, which can be experimentally realized by adding EGFR blocking antibodies to the system. We model the situation by introducing a scaling parameter η to the number of receptors, 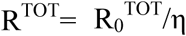, where subscript 0 indicates the reference values (Table 1 in the Supporting Material). Assuming the total number of EGFR becomes 20% of the original condition after antibody addition (η = 5), the two degrees of chemotaxis show different behaviors when compared to the unperturbed systems ∆R° (Fig. 3, magenta). The SNR-based 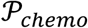 model predicts that antibody-treated cells become more sensitive than the control group at a low level of EGF (Fig. 3*B*), while the traditional ∆R° model does not show this nontrivial pattern (Fig. 3A). The intersecting point occurs at 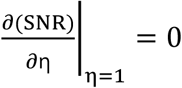, which can be translated into ~0.672ng/ml with the experimental values of this work (Supporting Material).

**FIGURE 3.**
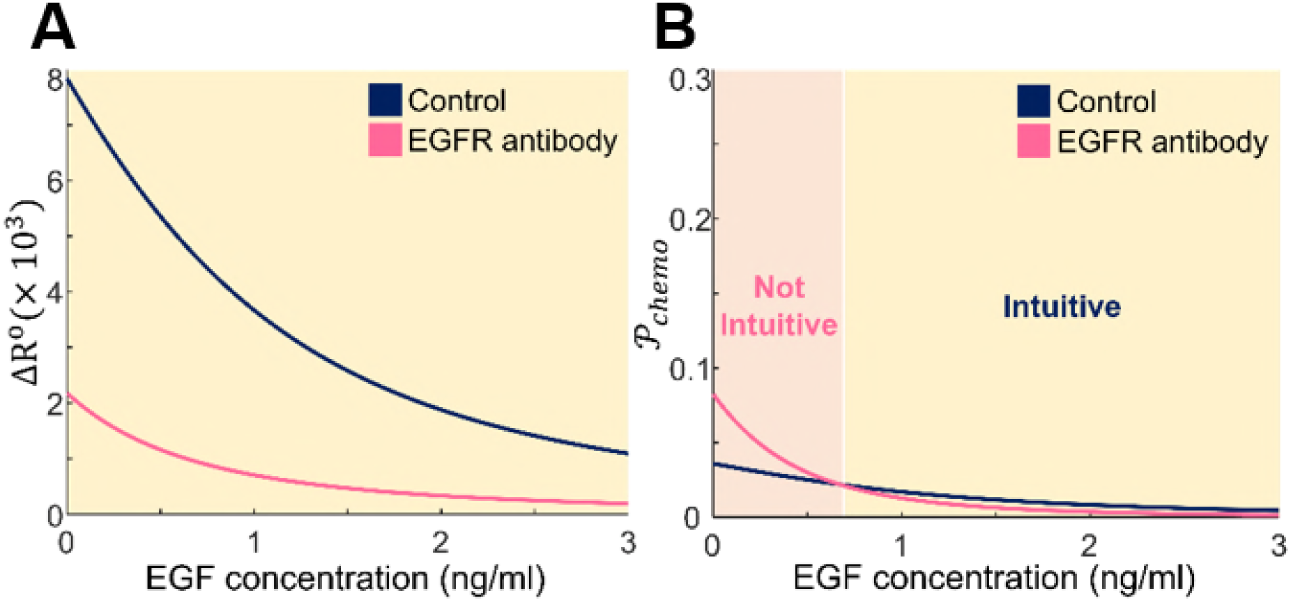
EGF concentration dependencies on two theoretical models of chemotaxis. (A) The difference of the number of bound receptors on anterior and posterior ends of a single cell, denoted by ∆R°, decreases as EGF concentration rises. Reducing the number of full-length EGFR through the addition of anti-EGFR antibody further reduces ∆R° compared to the control case. (B) The calculated propensity of chemotaxis, 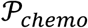, exhibits similar decreasing behavior to ∆R° with increasing EGF concentration, but in particular case with low EGF concentration, 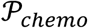 increased when the number of EGFR is reduced. The parameters used for theoretical models are listed in the Table S1. The dark blue curves correspond to the control group (η = 1), and magenta curves represent EGFR antibody treated group (η = 5).

### EGFR antibody can promote chemotaxis at a low level of EGF

As the two theoretical models predict different behaviors, we conducted another set of microfluidic experiments now with blocking antibodies to confirm which prediction is correct. We treated the cancer cells with two different concentrations (1 ng/ml and 10 μg/ml) of anti-EGFR antibodies and placed the cells under the EGF concentration gradient of a comparatively low slope (0 to 1 ng/ml). We also tested untreated cells for control, and other experimental settings remained the same. The chemotaxis behaviors are quantified as before in terms of the angle distribution of net cell migration and the CI. As shown in Fig. 4, the control group shows a pattern more similar to random motion, but the cells treated with 1 ng/ml antibody move with a significantly stronger bias in response to the chemical gradient. The CI for each group are −0.003 ± 0.041 (control, without antibody), 0.121 ± 0.037 (antibody of 1 ng/ml), and −0.008 ± 0.054 (antibody of 10 μg/ml), and the κ values are 0.0156 for the control group and 0.3965 for the group treated with 1 ng/ml blocking antibody. This finding is particularly interesting since it is apparently contradictory to the conventional notion that the reduction of EGFR always desensitizes cellular chemotaxis (34). On the other hand, the addition of high-dosage anti-EGFR antibody (10 μg/ml) reduced chemotaxis. The excessive reduction of EGFR may have mitigated the depletion of ligands, which presumably led to a decline in the magnitude of SNR. This experimental result suggests that the quantitative propensity of chemotaxis is a better description of how cells react to chemicals in a fluctuating environment than a traditional index of the number difference.

**FIGURE 4.**
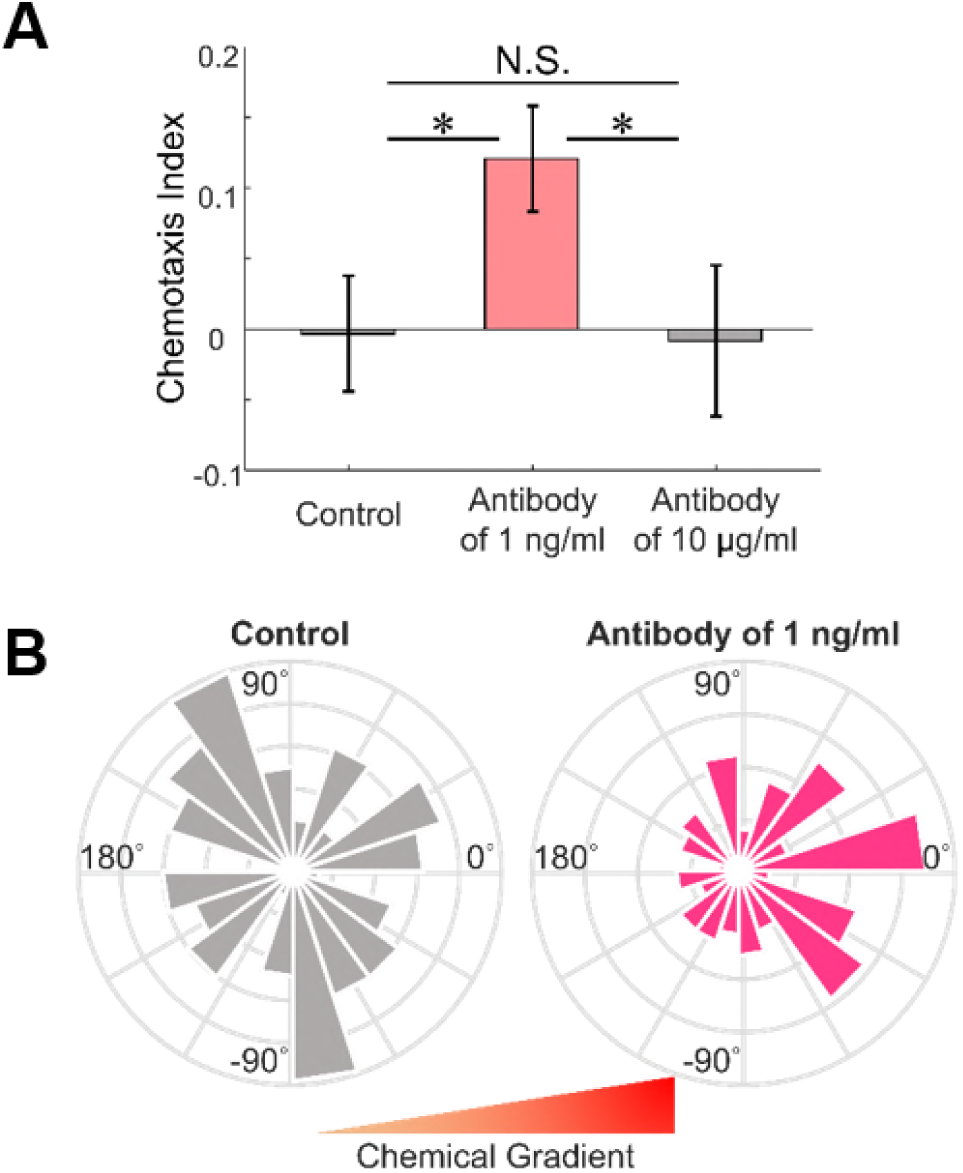
Effects of EGFR antibody on chemotaxis. (A) Comparison of CI and (B) angle histogram plots for the azimuth angle show that the control group represents a pattern more similar to random motion, but the cells treated with 1 ng/ml anti-EGFR antibody move with a significantly stronger bias in response to the chemical gradient. However, the addition of high-dosage antibody (10 μg/ml) reduces chemotaxis. Mean ± SEM (n = 123 for control, 151 for low-dosage, and 87 for high-dosage). * p<0.05.

## DISCUSSION

In this work, we combine theoretical and experimental approaches to show that local concentrations of the ligand affect chemotaxis behaviors, even when cancer cells are under the same linear concentration gradients. Moreover, we have demonstrated that reducing the number of full-length EGFR can raise the inclination of chemotaxis at low levels of EGF. These findings can be interpreted by our theoretical model for chemotaxis in terms of ligand perturbation and ligand depletion, and we attempt to address the importance of the two phenomena in certain conditions which leads to increase in chemotaxis. Finally, the results stress the need for quantitative analysis of chemotaxis when determining the dosage for anticancer therapy.

To test the robustness of our chemotaxis model to parameter perturbations, we conducted sensitivity analysis, where the concentration of EGF is fixed at 0.5 ng/ml, and the other reference parameters remain the same (Table 1 in the Supporting Material). We changed α (binding rate), β (dissociation rate), and D (diffusion coefficient) independently, by their relative magnitudes (to their reference values) from 1/2 to 2. Then we monitored 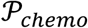 from the model to see to what extent each parameter affects the output. For a range of EGF concentrations (0 to 5 ng/ml), the response of 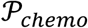 is plotted in Fig. S7 and the behaviors are consistent with the previous analysis. As shown in Fig. 5*A*, the chemotaxis propensity value remained in a reasonable realm in spite of the change of parameters by two folds change in both increasing and decreasing direction, indicating the robustness of our model. Among different parameters, the effects of the dissociation rate and the diffusion coefficient on 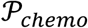 are more significant than the effect of the binding rate.

**FIGURE 5.**
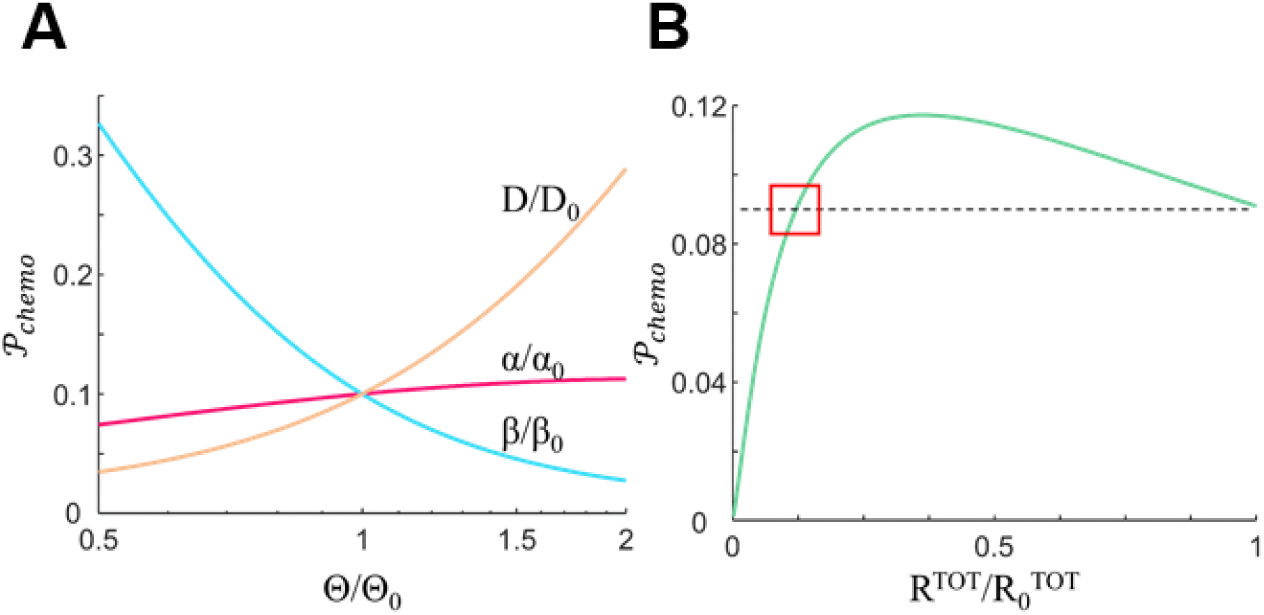
Sensitivity analysis of the chemotaxis. (A) The values of 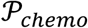 depend on relative variations of the parameters where α is the binding rate; β is the dissociation rate; D is the diffusion coefficient; subscript 0 indicates the reference values. The effects of the dissociation rate and the diffusion coefficient on 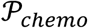 are more significant than the effect of binding rate. (B) The variations of 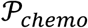 depending on the total number of receptors show that 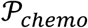 may increase upon reduction of EGFR. The green line corresponds to 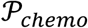, and the red square represents the critical point where the value of 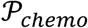 upon reduction of EGFR is equal to the value of 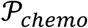 at the reference condition.

Furthermore, we noted that 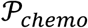 may increase upon reduction of EGFR (Fig. 5*B*) until the critical point as shown in Fig. 5*B* with the red square, where the value of 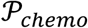 upon reduction of EGFR is equal to the value of 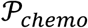 at the reference condition. Further reduction beyond the critical point may result in the gradual decrease in sensing ability, as depicted by low 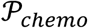. The partial reduction in the number of EGFR may have sensitized the cells in response to perturbing concentration, leading to increase in 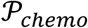. In low ligand concentration, as ligands bind to receptors, ligand concentration changes significantly and the value for the fraction of occupied receptors over total receptors also shifts, following the rules of chemical kinetics. The change in local ligand concentration causes change in fluctuations of concentrations, which affects 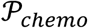. Shankaran et al. showed similar phenomenon for desensitization of G protein-coupled receptor (GPCR), based on a theoretical model. Desensitization of GPCR improves the information processing ability in response to ligand perturbations due to a reduced time delay in activation of receptor-ligand complexes (40), but the same desensitization also reduces GPCR signaling (41). β-arrestin, a GPCR binding factor, plays a similar role of an EGFR antibody in our model, and such approach of including activation step of a receptor-ligand complex could be further applied to improving our model.

We further note that in previous SNR-based chemotaxis studies, SNR profiles were reported as the bell shape (13, 16, 17), whereas our model predicts a unique monotonic behavior of 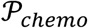 (Fig. 3*B*). The difference in profile is due to the differences in the chemical gradient in which chemotactic behaviors have taken place. Our result is obtained by employing a linear chemical gradient while the previous works used nonlinear chemical gradients where a chemical gradient is proportional to the chemical concentration. When our theoretical model is also applied to a nonlinear gradient, the profile of SNR results in the form of lagged bell shape curve along the local concentration, similar to previous results (Fig. S5). With the linear gradient, the effects of local concentration and steepness can be analyzed separately, thereby enabling the study of chemotaxis with different local concentration under the same linear gradient, as we have shown in Figs. 1 and 2. The linear gradient may also be pathologically significant since a chemical gradient by paracrine signaling between an immune cell and a cancer cell can be linear (42).

The cell type used in the experiment, MDA-MB-231, is a human metastatic breast cancer cell characterized by triple negative breast cancer (TNBC), which has insufficient biomarkers to be used for diagnosis and treatment (43-45). EGFR target therapy is an alternative strategy for TNBC to overcome the clinical challenges (45, 46). However, previous in vivo studies have shown that the EGF concentration is not very high in many parts of the body (47-49). Our finding that chemotaxis of breast cancer cells may be increased with reduced number of EGFR at a low EGF concentration hence implies the need for careful determination of the dosage for EGF target therapy to prevent possible unintentional aggravation.

## CONCLUSIONS

We modeled chemotaxis of breast cancer cells under EGF gradient by performing the theoretical analysis of SNR and conducting experiments at low EGF concentrations using a microfluidic system. As a result, we observed an interesting phenomenon that reducing the number of EGFR with the addition of anti-EGFR antibody can better direct cancer cell migration. The demonstration that external noise may change the chemotaxis pattern at low ligand concentration could provide clinically valuable insight into cancer therapy.

## AUTHOR CONTRIBUTIONS

S.L and J.S.J designed research. S.L. and H.N. fabricated microfluidic systems, and performed microfluidic experiments. S.L. obtained all meaurements and analyze the experimental data. S.L. developed an in-house code to simulate chemotaxis and a mathematical expression. The results were discussed critically with all authors, and S.L. and J.S.J. and wrote the manuscrpit. J.S.J. supervised this research.

## ACKNOWLEDGMENTS

The authors would like to acknowledge Jeong-Mo Choi for helpful discussion with simulation results and manuscript preparation. This research is supported by Basic Science Research Program through the National Research Foundation of Korea (NRF) funded by the Ministry of Education (2017R1D1A1B03030428) and the Brain Korea 21 Plus project.

## REFERENCES

1. Roussos, E. T., J. S. Condeelis, and A. Patsialou. 2011. Chemotaxis in cancer. Nat. Rev. Cancer 11:573–587.

2. Young, A., and C.-E. McNaught. 2011. The physiology of wound healing. Surgery (Oxford) 29:475–479.

3. Shahrara, S., S. R. Pickens, A. Dorfleutner, and R. M. Pope. 2009. IL-17 induces monocyte migration in rheumatoid arthritis. J. Immunol. 182:3884–3891.

4. Stokes, C. L., and D. A. Lauffenburger. 1991. Analysis of the roles of microvessel endothelial cell random motility and chemotaxis in angiogenesis. J. Theor. Biol. 152:377–403.

5. Müller, A., B. Homey, H. Soto, N. Ge, D. Catron, M. E. Buchanan, T. McClanahan, E. Murphy, W. Yuan, and S. N. Wagner. 2001. Involvement of chemokine receptors in breast cancer metastasis. Nature 410:50–56.

6. Joyce, J. A., and J. W. Pollard. 2009. Microenvironmental regulation of metastasis. Nat. Rev. Cancer 9:239–252.

7. Iglesias, P. A., and P. N. Devreotes. 2008. Navigating through models of chemotaxis. Curr. Opin. Cell Biol. 20:35–40.

8. Lauffenburger, D. A. 1982. Influence of external concentration fluctuations on leukocyte chemotactic orientation. Cell Biophys. 4:177.

9. Tranquillo, R., and D. Lauffenburger. 1988. Analysis of leukocyte chemosensory movement. Adv Biosci 66:29–38.

10. Tranquillo, R. T., D. A. Lauffenburger, and S. Zigmond. 1988. A stochastic model for leukocyte random motility and chemotaxis based on receptor binding fluctuations. J. Cell. Biol. 106:303–309.

11. Tranquillo, R. T., and D. A. Lauffenburger. 1987. Stochastic model of leukocyte chemosensory movement. J. Math. Biol. 25:229–262.

12. Shibata, T., and K. Fujimoto. 2005. Noisy signal amplification in ultrasensitive signal transduction. Proc. Natl. Acad. Sci. U.S.A. 102:331–336.

13. Ueda, M., and T. Shibata. 2007. Stochastic signal processing and transduction in chemotactic response of eukaryotic cells. Biophys. J. 93:11–20.

14. Van Haastert, P. J., and M. Postma. 2007. Biased random walk by stochastic fluctuations of chemoattractant-receptor interactions at the lower limit of detection. Biophys. J. 93:1787–1796.

15. Levine, H., and W. J. Rappel. 2013. The physics of eukaryotic chemotaxis. Phys. Today 66.

16. Mortimer, D., J. Feldner, T. Vaughan, I. Vetter, Z. Pujic, W. J. Rosoff, K. Burrage, P. Dayan, L. J. Richards, and G. J. Goodhill. 2009. A Bayesian model predicts the response of axons to molecular gradients. Proc. Natl. Acad. Sci. U.S.A. 106:10296–10301.

17. Mortimer, D., Z. Pujic, T. Vaughan, A. W. Thompson, J. Feldner, I. Vetter, and G. J. Goodhill. 2010. Axon guidance by growth-rate modulation. Proc. Natl. Acad. Sci. U.S.A. 107:5202–5207.

18. Fuller, D., W. Chen, M. Adler, A. Groisman, H. Levine, W. J. Rappel, and W. F. Loomis. 2010. External and internal constraints on eukaryotic chemotaxis. Proc. Natl. Acad. Sci. U.S.A. 107:9656–9659.

19. Neumann, S., L. Lovdok, K. Bentele, J. Meisig, E. Ullner, F. S. Paldy, V. Sourjik, and M. Kollmann. 2014. Exponential signaling gain at the receptor level enhances signal-to-noise ratio in bacterial chemotaxis. PLoS One 9:e87815.

20. Amselem, G., M. Theves, A. Bae, C. Beta, and E. Bodenschatz. 2012. Control parameter description of eukaryotic chemotaxis. Phys Rev Lett 109:108103.

21. Segota, I., and C. Franck. 2017. Extracellular Processing of Molecular Gradients by Eukaryotic Cells Can Improve Gradient Detection Accuracy. Phys. Rev. Lett. 119:248101.

22. Friedl, P., and K. Wolf. 2009. Plasticity of cell migration: a multiscale tuning model. J. Cell Biol.:jcb. 200909003.

23. Zigmond, S. H. 1977. Ability of polymorphonuclear leukocytes to orient in gradients of chemotactic factors. J. Cell Biol. 75:606–616.

24. Zicha, D., G. A. Dunn, and A. F. Brown. 1991. A new direct-viewing chemotaxis chamber. J. Cell Sci. 99:769–775.

25. Jeon, N. L., H. Baskaran, S. K. Dertinger, G. M. Whitesides, L. Van De Water, and M. Toner. 2002. Neutrophil chemotaxis in linear and complex gradients of interleukin-8 formed in a microfabricated device. Nat. Biotechnol. 20:826–830.

26. Knapp, D. M., E. F. Helou, and R. T. Tranquillo. 1999. A fibrin or collagen gel assay for tissue cell chemotaxis: assessment of fibroblast chemotaxis to GRGDSP. Exp. Cell Res. 247:543–553.

27. Rosoff, W. J., J. S. Urbach, M. A. Esrick, R. G. McAllister, L. J. Richards, and G. J. Goodhill. 2004. A new chemotaxis assay shows the extreme sensitivity of axons to molecular gradients. Nat. Neurosci. 7:678–682.

28. Lauffenburger, D. A., and S. H. Zigmond. 1981. Chemotactic factor concentration gradients in chemotaxis assay systems. J. Immunol. Methods 40:45–60.

29. Polacheck, W. J., J. L. Charest, and R. D. Kamm. 2011. Interstitial flow influences direction of tumor cell migration through competing mechanisms. Proc. Natl. Acad. Sci. U.S.A. 108:11115–11120.

30. Jeon, J. S., S. Bersini, M. Gilardi, G. Dubini, J. L. Charest, M. Moretti, and R. D. Kamm. 2015. Human 3D vascularized organotypic microfluidic assays to study breast cancer cell extravasation. Proc. Natl. Acad. Sci. U.S.A. 112:214–219.

31. Ramanujan, S., A. Pluen, T. D. McKee, E. B. Brown, Y. Boucher, and R. K. Jain. 2002. Diffusion and convection in collagen gels: implications for transport in the tumor interstitium. Biophys. J. 83:1650–1660.

32. Martino, M. M., P. S. Briquez, A. Ranga, M. P. Lutolf, and J. A. Hubbell. 2013. Heparin-binding domain of fibrin (ogen) binds growth factors and promotes tissue repair when incorporated within a synthetic matrix. Proc. Natl. Acad. Sci. U.S.A. 110:4563–4568.

33. Wang, S.-J., W. Saadi, F. Lin, C. M.-C. Nguyen, and N. L. Jeon. 2004. Differential effects of EGF gradient profiles on MDA-MB-231 breast cancer cell chemotaxis. Exp. Cell Res. 300:180–189.

34. Saadi, W., S.-J. Wang, F. Lin, and N. L. Jeon. 2006. A parallel-gradient microfluidic chamber for quantitative analysis of breast cancer cell chemotaxis. Biomed. Microdevices 8:109–118.

35. Szatmary, A. C., and R. Nossal. 2017. Determining whether observed eukaryotic cell migration indicates chemotactic responsiveness or random chemokinetic motion. J. Theor. Biol. 425:103–112.

36. Shin, Y., S. Han, J. S. Jeon, K. Yamamoto, I. K. Zervantonakis, R. Sudo, R. D. Kamm, and S. Chung. 2012. Microfluidic assay for simultaneous culture of multiple cell types on surfaces or within hydrogels. Nat. Protoc. 7:1247–1259.

37. Piccinini, F., A. Kiss, and P. Horvath. 2015. CellTracker (not only) for dummies. Bioinformatics 32:955–957.

38. Berens, P. 2009. CircStat: a MATLAB toolbox for circular statistics. J. Stat. Softw. 31:1–21.

39. Liu, H., Y.-D. Cao, W.-X. Ye, and Y.-Y. Sun. 2010. Effect of microRNA-206 on cytoskeleton remodelling by downregulating Cdc42 in MDA-MB-231 cells. Tumori 96:751–755.

40. Shankaran, H., H. S. Wiley, and H. Resat. 2007. Receptor downregulation and desensitization enhance the information processing ability of signalling receptors. BMC Syst. Biol. 1:48.

41. Rajagopal, S., and S. K. Shenoy. 2017. GPCR desensitization: acute and prolonged phases. Cell. Signal.

42. Kim, B. J., and M. Wu. 2012. Microfluidics for mammalian cell chemotaxis. Annals of biomedical engineering 40:1316–1327.

43. Foulkes, W. D., I. E. Smith, and J. S. Reis-Filho. 2010. Triple-negative breast cancer. N. Engl. J. Med. 363:1938–1948.

44. Bianchini, G., J. M. Balko, I. A. Mayer, M. E. Sanders, and L. Gianni. 2016. Triple-negative breast cancer: challenges and opportunities of a heterogeneous disease. Nat. Rev. Clin. Oncol. 13:674–690.

45. Hudis, C. A., and L. Gianni. 2011. Triple-negative breast cancer: an unmet medical need. Oncologist 16 Suppl 1:1–11.

46. Ferraro, D. A., N. Gaborit, R. Maron, H. Cohen-Dvashi, Z. Porat, F. Pareja, S. Lavi, M. Lindzen, N. Ben-Chetrit, M. Sela, and Y. Yarden. 2013. Inhibition of triple-negative breast cancer models by combinations of antibodies to EGFR. Proc. Natl. Acad. Sci. U.S.A. 110:1815–1820.

47. Playford, R., and S. Ghosh. 2005. Cytokines and growth factor modulators in intestinal inflammation and repair. J. Pathol. 205:417–425.

48. Brand, H. S., A. J. Ligtenberg, and E. C. Veerman. 2014. Saliva and wound healing. Monogr. Oral Sci. 24:52–60.

49. Kazemnejad, S., A. Allameh, A. Gharehbaghian, M. Soleimani, N. Amirizadeh, and M. Jazayeri. 2008. Efficient replacing of fetal bovine serum with human platelet releasate during propagation and differentiation of human bone marrow-derived mesenchymal stem cells to functional hepatocytes-like cells. Vox Sang. 95:149–158.

50. Fitzpatrick SL, LaChance MP, Schultz GS. Characterization of epidermal growth factor receptor and action on human breast cancer cells in culture. Cancer Res. 44, 3442–3447 (1984).

51. Lang NR, et al. Estimating the 3D pore size distribution of biopolymer networks from directionally biased data. Biophys. J. 105, 1967–1975 (2013).

52. T horne RG, Hrabetova S, Nicholson C. Diffusion of epidermal growth factor in rat brain extracellular space measured by integrative optical imaging. J. Neurophysiol. 92, 3471–3481 (2004).

53. Bailly M, et al. Epidermal growth factor receptor distribution during chemotactic responses. Mol. Biol. Cell 11, 3873–3883 (2000).

54. Hughes B. On the error probability of signals in additive white Gaussian noise. IEEE Trans. Inf. Theory 37, 151–155 (1991).

55. Schoeberl B, Eichler-Jonsson C, Gilles ED, Müller G. Computational modeling of the dynamics of the MAP kinase cascade activated by surface and internalized EGF receptors. Nat. Biotechnol. 20, 370–375 (2002).

56. Zhou M, et al. Real-time measurements of kinetics of EGF binding to soluble EGF receptor monomers and dimers support the dimerization model for receptor activation. Biochemistry 32, 8193–8198 (1993).

